# Biosynthesis of Steroidal Alkaloids Are Coordinately Regulated and Differ Among Tomatoes in the Red-Fruited Clade

**DOI:** 10.1101/2021.01.06.425594

**Authors:** Michael P. Dzakovich, David M. Francis, Jessica L. Cooperstone

## Abstract

- We quantitatively profiled and genotyped two tomato populations representing diversity in the red-fruited clade to address the lack of knowledge regarding the chemical diversity, concentration, and genetic architecture controlling tomato steroidal alkaloids.
- We grew 107 genetically diverse fresh market, processing, land-race, and wild tomatoes in multiple environments. Nine steroidal alkaloids were quantified using ultra-high performance liquid chromatography tandem mass spectrometry. The diversity panel and a biparental population segregating for high alpha-tomatine, were genotyped to identify and validate quantitative trait loci (QTL) associated with steroidal alkaloids.
- Land-races and wild material exhibited higher alkaloid concentrations and more chemical diversity. Average total content of steroidal alkaloids, often dominated by lycoperoside F/G/esculeoside A, ranged from 1.9 to 23.3 mg/100 g fresh weight across accessions. Land-race and wild cherry accessions distinctly clustered based on elevated early or late steroidal alkaloid concentrations. Significant correlations were observed among early and late steroidal alkaloids in a species-dependent manner. A QTL controlling multiple, early steroidal alkaloid pathway intermediates on chromosome 3 was identified by genome wide association (GWAS) and validated in a backcross population.
- Tomato steroidal alkaloids are diverse in the red-fruited tomato clade and their biosynthesis is regulated in a coordinated manner.

## 1.2 Introduction

The tomato clade of Solanaceae produce a unique assortment of cholesterol-derived steroidal alkaloids (Cárdenas *et al*., 2015). These secondary metabolites are thought to be produced for plant defense, due to their reported fungicidal and insecticidal properties (Irving *et al*., 1945; Cipollini & Levey, 1997). Alpha-tomatine, the most well studied tomato steroidal alkaloid, was hypothesized to impart resistance to *Fusarium* in the mid 20^th^ century (Gottlieb, 1943; Irving *et al*., 1945) and was isolated and characterized shortly thereafter (Fontaine *et al*., 1948). While steroidal alkaloids are frequently considered anti-nutritional compounds (Itkin *et al*., 2013; Cárdenas *et al*., 2016; Ballester *et al*., 2016), a growing body of literature suggests that tomato steroidal alkaloids may impart positive health benefits as part of the diet (Friedman *et al*., 2000; Lee *et al*., 2004; Choi *et al*., 2012a; Dyle *et al*., 2014).

Researchers have used association analysis, comparative transcriptomics, and genetic modification as well as high resolution mass spectrometry to describe the steroidal alkaloid biosynthetic pathway and its chemical constituents (Mintz-Oron *et al*., 2008; Itkin *et al*., 2011; Iijima *et al*., 2013; Schwahn *et al*., 2014; Alseekh *et al*., 2015; Ballester *et al*., 2016; Abdelkareem *et al*., 2017; Sonawane *et al*., 2018; Zhu *et al*., 2018; Bednarz *et al*., 2019; Cárdenas *et al*., 2019). These recent efforts to elucidate the steroidal alkaloid biosynthetic pathway and its regulatory mechanisms in tomato are bearing fruit (Itkin *et al*., 2011, 2013; Ballester *et al*., 2016; Sonawane *et al*., 2018; Cárdenas *et al*., 2019; Yu *et al*., 2020). GLYCOALKALOID METABOLISM (GAME) enzymes 4, 6, 7, 8, 11, and 12 have been shown to catalyze a series of hydroxylation, oxidation, and transamination reactions on the aliphatic tail of cholesterol to generate the E and nitrogenous F rings characteristic of solanaceous steroidal alkaloids (Itkin *et al*., 2013). This series of reactions results in the biosynthesis of dehydrotomatidine, the first steroidal alkaloid in the proposed pathway. Dehydrotomatidine can be converted into tomatidine by GAME25, SlS5αR2 (a C5-alpha reductase), and Sl3βHSD1 (a C3-dehydrogenase/reductase) (Sonawane *et al*., 2018; Akiyama *et al*., 2019; Lee *et al*., 2019). It is been proposed that the pathway then branches into desaturated (derived from dehydrotomatidine) and saturated (derived from tomatidine) steroidal alkaloids which are biosynthesized in parallel using the same enzymes at each step. Dehydrotomatidine and tomatidine are then converted to dehydrotomatine and alpha-tomatine, respectively, by a series of glycosylations catalyzed by GAME1, 17, 18, and 2 (Itkin *et al*., 2011, 2013). These compounds can then be hydroxylated into hydroxy-dehydrotomatine and hydroxytomatine by 2-oxoglutarate-dependent dioxygenases (GAME31 and 32) (Cárdenas *et al*., 2019). Based on chemical structure the next steps are presumed to follow the order hydroxytomatine, acetoxytomatine, lycoperoside F/G/esculeoside A, and esculeoside B (and their desaturated counterparts). Recent evidence indicates that GAME5 (a uridine diphosphate glycosyltransferase) is critical for the enzymatic conversion of acetoxytomatine to esculeoside A (Szymański *et al*., 2020). However, the structural and regulatory mechanisms responsible for the production of other steroidal alkaloids occurring in the later part of the pathway have yet to be described.

Alpha-tomatine is the primary steroidal alkaloid in mature green tomatoes and is converted into various downstream lycoperosides and esculeosides as the fruit matures (Iijima *et al*., 2009; Cárdenas *et al*., 2015). Natural variation in this process has been observed in a small subset of *Solanum lycopersicum* var. *cerasiforme* Latin American cultivar accessions, which may be considered land-races, including LA2213, LA2256, and LA2262 (Rick *et al*., 1994). Fruits from these accessions retain high levels of alpha-tomatine throughout ripening contrary to most members of the red-fruited clade, and are commonly consumed in the Alto Mayo region of Peru (Rick *et al*., 1994). These accessions, along with other selections made based on known patterns of genetic diversity, provide variability that may offer insight into the regulation of the tomato steroidal alkaloid biosynthetic pathway in the red-fruited clade.

The purpose of our work is to describe the chemical diversity of tomato steroidal alkaloids through a detailed and quantitative analysis (Dzakovich *et al*., 2020) of cultivated, land-race, and wild red-fruited relatives of tomato and identify quantitative trait loci (QTL) associated with these metabolites. Populations were selected based on known patterns of genomic variation and were cultivated in multiple locations and years to separate genetic and environmental effects. We quantified tomato steroidal alkaloids in a diversity panel of 107 red-fruit tomato accessions and determined their associated QTL by a genome-wide association analysis (GWAS). Finally, we estimated genetic effects and validated findings from the GWAS by generating, genotyping and phenotyping a bi-parental population based on high alpha-tomatine germplasm crossed to a cultivated parent.

## 1.3 Materials and Methods

### 1.3.1 Plant Material

A panel of 107 tomato accessions from public breeding programs, the C.M. Rick Tomato Genetics Resource Center (TGRC), and the United States Department of Agriculture (USDA) National Plant Germplasm System (NPGS) was assembled (Table **S1**). Selection criteria included known patterns of genetic diversity as determined by genotyping using a 7,720 SNP array (Sim *et al*., 2012a; Blanca *et al*., 2015), previous phenotypic information for alkaloid content (Rick *et al*., 1994), and geographic origin (visualized in Fig. **S1**.). To maximize genetic diversity within the red-fruited tomato species, selections included *S. pimpinellifolium, S. lycopersicum* var. *cerasiforme* (including wild, land-race, and cultivated cherry tomatoes), and *S. lycopersicum*. Sub-population sizes were chosen based on rarefaction analysis to capture at least of 85% of the genetic variation in previously defined sub-populations (Blanca *et al*., 2015). This approach resulted in three species with 32 sub-populations of previously classified material: 25 accessions of *S*.*pimpinellifolium* (representing 11 sub-populations), 37 accessions of *S. lycopersicum* var. *cerasiforme* (representing 14 sub-populations; including six cultivated cherry varieties), and 39 accessions of *S. lycopersicum* (representing 7 sub-populations) (Blanca *et al*., 2015). The *S. lycopersicum* selections included processing varieties representing the diversity present in this market class (Sim *et al*., 2012a; Blanca *et al*., 2015). Heinz 1706 (LA4345) and M82 (LA3475) were included as part of the cultivated processing tomatoes to serve as standard reference material due to their widespread and historical use in plant biology research. For subsequent analyses, we divide this population into five groups: cultivated processing, cultivated cherry, wide-cross hybrid, wild cherry (including land-races), and *S. pimpinellifolium* (Table **S1**). A list of all the material used in this diversity panel can be found in the supporting information (Table **S1**).

### 1.3.2 Field Trial

Plants were grown in three field environments during the summers of 2017 and 2018 at the North Central Agricultural Research Station in Fremont, OH. Summer, 2017 included the first field environment (planted 5/24/2017) while the remaining two (early planting (5/29/2018); late planting (6/18/2018)) were grown during 2018. Environments are differentiated by time (year and/or planting date) and geophysical conditions such as soil composition. In all three environments, plants were grown in a randomized complete block design with two blocks per environment (total n=642). Within each environment, plants were grown in plots comprised of 6-10 plants and samples were an aggregate of fruits from all plants within a plot excluding individuals at the ends of each plot. Red ripe fruits were harvested over a three-week period from each environment and stored whole at −40 ° C until analysis.

### 1.3.3 Inbred Backcross Population Development

Three accessions of tomatoes containing high levels of alpha-tomatine in red-ripe fruits (LA2213, LA2256, and LA2262) as well as the cultivated processing accession OH8243 (PI 601423) and the cultivated cherry accession Tainan (PI 647556) variety were also cultivated under glasshouse conditions during Spring 2017 and crossed. Seedlings from the resulting six sets of F_1_ progeny as well as parental material were included in the 2017 and 2018 field trials and profiled for steroidal alkaloids as described below. F_1_ individuals from OH8243×LA2213 were backcrossed to the cultivated processing tomato recurrent parent (OH8243) and 200 individual BC1 plants were grown under glasshouse conditions during Spring 2018. Progeny were then field grown during Summer 2018, and BC_1_S_1_ individuals were genotyped and profiled for steroidal alkaloids as described below.

### 1.3.4 Steroidal Alkaloid Profiling

Steroidal alkaloids were extracted and quantified, as previously described, using a rapid extraction and ultra-high performance liquid chromatography tandem mass spectrometry (UHPLC-MS/MS) (Dzakovich *et al*., 2020). Dehydrotomatidine, tomatidine, dehydrotomatine, alpha-tomatine, hydroxytomatine, acetoxytomatine, dehydrolycoperoside F/G/dehydroesculeoside A, lycoperoside F/G/esculeoside A, esculeoside B, and total steroidal alkaloids (the sum of all previous analytes) were quantified. Alkaloids that have the same mass (e.g., lycoperoside F/G/esculeoside A) were summer together if they could not be differentiated (Dzakovich *et al*., 2020). Acetonitrile, formic acid, isopropanol, methanol, and water were of LC-MS grade and purchased from Fisher Scientific (Pittsburgh, PA). Alpha-tomatine (≥90% purity) and solanidine (≥99% purity) were purchased from Extrasynthese (Genay, France). Alpha-solanine (≥95% purity) and tomatidine (≥95% purity) were purchased from Sigma Aldrich (St. Louis, MO).

### 1.3.5 Genotyping

DNA was extracted using a hexadecyltrimethylammonium bromide (CTAB) procedure optimized for tomato and scaled to accommodate 96-tube racks (Sim *et al*., 2015). Leaflet samples were homogenized with 60 μL of 5% Sarkosyl, 150 μL of CTAB lysis buffer (2 M NaCl, 0.2 M Tris, 0.05 M EDTA, and 2% CTAB maintained at a 7.5 pH), and 150 μL of extraction buffer (0.35 M sorbitol, 0.1 M Tris, 25 mM sodium bisulfite, and 5 mM EDTA maintained at pH 7.5) in 1.2 mL tubes. Two 4 mm metal beads were added to each tube prior to sealing and the rack was shaken at 200 rpm in a GenoGrinder (OPS Diagnostics, Lebanon, NJ) for two min. Racks were then placed in a 65 °C incubator for 20 min and then left to cool at room temperature for an additional 10 min. Each tube was then spiked with 350 μL of chloroform: isoamyl alcohol (24:1) and phase separation was induced by centrifuging plates for 10 min at 5,000 x *g*. Aqueous phases from each sample were transferred to a corresponding well on a fresh 96 well plate and DNA was precipitated by adding 110 μL of isopropyl alcohol to each well and centrifuging for an additional 15 min at 5000 x *g*. Plates were then uncovered and left to air dry for 30 min and DNA was resuspended in 100 μL of Tris-EDTA buffer (10 mM Tris and 0.1 mM EDTA).

Eighty five members of the diversity panel were previously genotyped for 7,720 SNPs using the SolCAP Illumina Infinium Array (Hamilton *et al*., 2012; Sim *et al*., 2012b,a; Blanca *et al*., 2015). To genotype the BC_1_S_1_ population, a 384 single nucleotide polymorphism (SNP) marker panel created by the Solanaceae Coordinated Agricultural Project (SolCAP) was utilized (Sim *et al*., 2015; Gill *et al*., 2019). Genetic positions based on Sim and colleagues (Sim *et al*., 2012b) and established patterns of linkage disequilibrium decay in *S. lycopersicum* (Robbins *et al*., 2011) were used to select SNPs distributed across all 12 chromosomes in regions of high recombination. Gaps in genome coverage were filled in by selecting high PIC markers based on physical position. Genotyping was conducted on 156 BC_1_S_1_ individuals using the PlexSeq™ platform (Agriplex Genomics, Cleveland, OH).

### 1.3.6 Data Analysis

#### 1.3.6.1 Linear modeling, heritability, and reliability

All statistical analyses and data visualization were conducted using R version 3.5.1 (R Development Core Team, 2018). A linear model based on our experimental design was used to estimate the effects of genetics and environment on steroidal alkaloid concentrations:

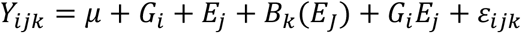

In the model above, *Y_ijk_* represents the estimate for a given analyte, *μ* represents the population mean of a given analyte, *G_i_* represents the contribution due to genetic factors, *E_j_* represents the contribution due to environmental factors, *B_k_*(*E_J_*) represents within environment effects, *G_i_E_j_* represents the interaction between genetic and environmental factors, and *ε_ijk_* represents the residual error. Variance components were estimated using “ranef” function in the package lme4 (Bates *et al*., 2015). Broad sense heritability (H^2^) was calculated according to previously described methods (Cotterill, 1987):

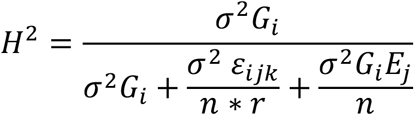

Where *σ*^2^ represents the variance, *n* is equal to the number of environments (3), *r* is the number of reps within an environment (2), and all other variables are as described above. Reliability (i^2^) was calculated with the previously defined model (Bernardo, 2020):

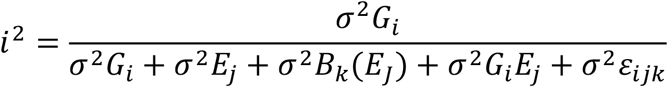

#### 1.3.6.2 GWAS

Association analysis was conducted using the R package rrBLUP with GWAS utilities (Endelman, 2011). SNP calls were converted to numerical scoring where 1 was homozygous for the common allele of the reference variety OH8245 (Berry *et al*., 1991). Homozygous for the alternative allele were scored as −1 and heterozygous calls were scored as 0. We used the unified mixed model to test for marker-trait associations (Yu *et al*., 2006). This model contains a matrix for kinship (K), considered a random effect, and fixed effects for structure (Q). The K matrix was obtained from the “a.mat” function in rrBLUP. The Q matrix was developed from principal components analysis (PCA), as described (Price *et al*., 2006). Briefly, a covariate matrix was calculated from the standardized n x M (number of lines x number of markers) matrix (N) by multiplying it by its transformation (N x N^−1^). The resulting n x n covariate matrix was then used as an input for PCA. We examined the effects of using 1 to 3 principal components (PC) by comparing Bayesian information criterion (BIC) for marker-trait models with different numbers of PC. For example, the naïve model *Y* = *Marker* (*M*) was compared to *Y* = *PC*1 + *M* and similarly the model with PC1 was compared to *Y* = *PC*1 + *PC*2 + *M* and *Y* = *PC*1 + *PC*2 + *PC*3 + *M* to test the effect of adding additional PC to the model. Benjamini-Hochberg corrected P values (Benjamini & Hochberg, 1995) were generated to control the overall false discovery rate (FDR; P < 0.05), account for multiple testing and provide balanced protection against type-I and -II error using the function “p.adjust.” The results of genome wide association analysis with QK models were visualized by plotting −log_10_ FDR corrected P-value vs genome position using ggplot2 (Wickham, 2016). The minor allele frequency (MAF) was set to 10% with alleles below the MAF threshold disregarded.

#### 1.3.6.3 QTL Analysis

Analysis of BC_1_S_1_ progeny were analyzed using ANOVA with the following model:

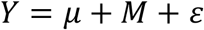

Where *Y* is the vector of steroidal alkaloid concentrations and *M* represents the effect of a given marker. Significant marker-trait associations were defined at a FDR-corrected P < 0.05 (Benjamini & Hochberg, 1995). P-values, the proportion of variance explained (r^2^), and the effects of allele substitution were summarized as a description of the QTL identified.

#### 1.3.6.4 Visualization of Steroidal Glycoalkaloid Variation

Least squares means were extracted from the fixed effects linear models for steroidal alkaloid concentrations for every genotype and used to generate covariance matrices used for principal components analysis (PCA) and correlation analyses (Table **S3**). Population structure and phenotypic variation was visualized using PCA with the packages FactoMineR and Factoextra using a correlation matrix with each variable scaled to a mean of zero and standard deviation of one as the input data (Lê *et al*., 2008). Boxplots and maps displaying the geographic origin of germplasm studied were generated using the packages ggplot2, ggmap, and maptools (Wickham, 2009; Kahle & Wickham, 2013; Bivand & Lewin-Koh, 2018). Correlations among analytes were visualized using the package corrplot (Wei & Simko, 2017).

## 1.4 Results

### 1.4.1 Land-races and wild tomato species exhibit steroidal alkaloid chemical diversity

We applied a recently developed method that extracts and quantifies 9 members of the proposed tomato steroidal alkaloid pathway (Dzakovich *et al*., 2020) to a population of 107 (Table **S1**) tomato accessions grown in three environments. We defined early and late pathway intermediates based on their position in the proposed steroidal alkaloid biosynthesis pathway, with dehydrotomatidine, tomatidine, dehydrotomatine, alpha-tomatine, hydroxytomatine and acetoxytomatine considered early and dehydrolycoperoside F/G/dehydroesculeoside A, lycoperoside F/G/esculeoside A, and esculeoside B as late. This definition of early and late pathway intermediates is reinforced by clustering patterns and correlation based on concentrations observed within tomato germplasm.

In the case of hydroxytomatine, acetoxytomatine, dehydrolycoperoside F/G/dehydroesculeoside A, lycoperoside F/G/esculeoside A, and esculeoside B, multiple peaks are observed, confirming the presence of structural isomers (Manabe *et al*., 2013; Nohara *et al*., 2015; Hövelmann *et al*., 2019; Steinert *et al*., 2020). For example, lycoperoside F, G, and esculeoside A (C58H95NO29, monoisotopic mass = 1269.5990 Da), are isobaric (Yahara *et al*., 2004) and exhibit similar fragmentation patterns since their molecular differences are limited to subtle changes in stereochemistry on the F ring (Dzakovich *et al*., 2020). Therefore, values reported here are the sum of all isomers within a given mass, and compounds are annotated as precisely as their identities are known. We report the sum of all steroidal alkaloids in a sample as means plus or minus standard deviation for each alkaloid. Genotype means and standard deviations for all steroidal alkaloids and their individual isomers can be found in the supporting information (Table **S2**). Differences of up to four orders of magnitude in steroidal alkaloid concentration were observed across the accessions selected. Box and whisker plots with log_10_ scaled y-axes detailing concentrations of each tomato steroidal alkaloid separated into five classes of tomato germplasm present in this collection can be found in Fig. **1**. Each dot within the box and whisker plots represents an individual observation. Means for all alkaloids were higher in land-races and wild species accessions when compared to cultivated material. A multi-modal distribution can be observed for concentrations of lycoperoside F/G/esculeoside A within the wild cherry tomato accession group (which also contains land-races). For all steroidal alkaloids measured, concentrations in cultivated material were lower relative to land-race and wild accessions.

**Fig. 1.**
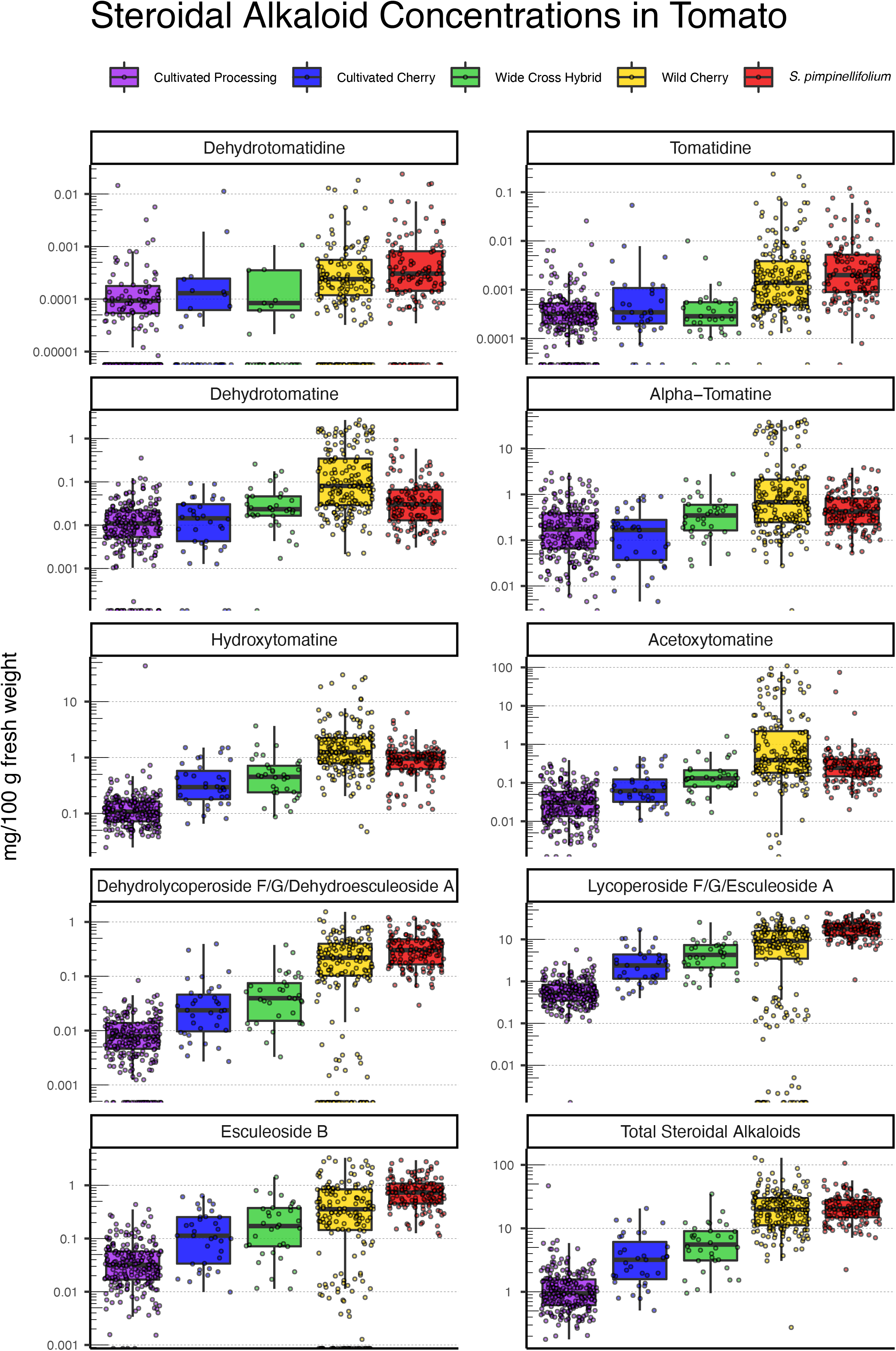
Box and whisker plots of steroidal alkaloid concentrations across all germplasm, separated by class. Each dot represents an individual observation. The y-axis was log transformed to visually condense the large amount of variation observed in the concentrations of all tomato steroidal alkaloids measured in this study. Distinct sub-groups of wild cherry tomatoes can be observed for lycoperoside F/G/esculeoside A

The predominant steroidal alkaloid in 85 of the 107 accessions was lycoperoside F/G/esculeoside A, a late pathway intermediate. Exceptions to this trend included high alpha-tomatine, hydroxytomatine, or acetoxytomatine land-race or wild accessions (Table **S1,2**). Lycoperoside F/G/esculeoside A comprised 82% of the total steroidal alkaloids in *S. pimpinellifolium* accessions, with similar proportions in the wide-cross hybrids (72%) and cultivated cherry (72%). The steroidal alkaloid profile of land-race and wild cherry and cultivated processing tomatoes was comprised of 37% and 48% lycoperoside F/G/esculeoside A, respectively. In land-race and wild cherry tomatoes, acetoxytomatine was the second most abundant steroidal alkaloid (28%) on average. Alpha-tomatine was the second most abundant (23%) steroidal alkaloid in cultivated processing tomatoes. However, alpha-tomatine only comprised 3%, 5%, and 20% of total steroidal alkaloids in *S. pimpinellifolium*, cultivated cherry, and wild cherry accessions. These generalities are not always true for each individual accession given the diversity that exists within each tomato class. Median concentrations of individual steroidal alkaloids were between 0.1 μg/100g and 1 mg/100 g fresh weight (Fig. **1**). For total steroidal alkaloids, median values were between 1 and 5 mg/100g fresh weight for cultivated processing and cultivated cherry, respectively. *S. pimpinellifolium*, land-race, and wild cherry had median total steroidal alkaloid values around 23 mg/100g fresh weight.

### 1.4.2 Early or late pathway steroidal alkaloids drive separation in land-race and wild cherry tomatoes

Differences in the overall profile of steroidal alkaloids from ripe fruit in accessions from the diversity panel were visualized using principal components analysis (PCA, Fig. **2**). Least squares means were calculated for each genotype, and these data were used as inputs for the PCA (Table **S3**). The first PC explained 30.1% of the phenotypic variance and the second PC, 26.8%. Cultivated material clusters strongly together. Two sub-groups of wild cherry fruits are apparent, both of which separate from cultivated cherry (Fig. **2A**). Hierarchical clustering using Ward’s D2 minimum variance linkage method confirmed that these individuals form unique clusters (Fig. **S2**). A loadings plot (Fig. **2B**) of the PCA indicates separation along PC1 is driven by total steroidal alkaloid content. Location in the tomato steroidal alkaloid biosynthesis pathway drives separation along PC2, with early pathway intermediates towards the bottom, and later pathway products at the top.

**Fig. 2.**
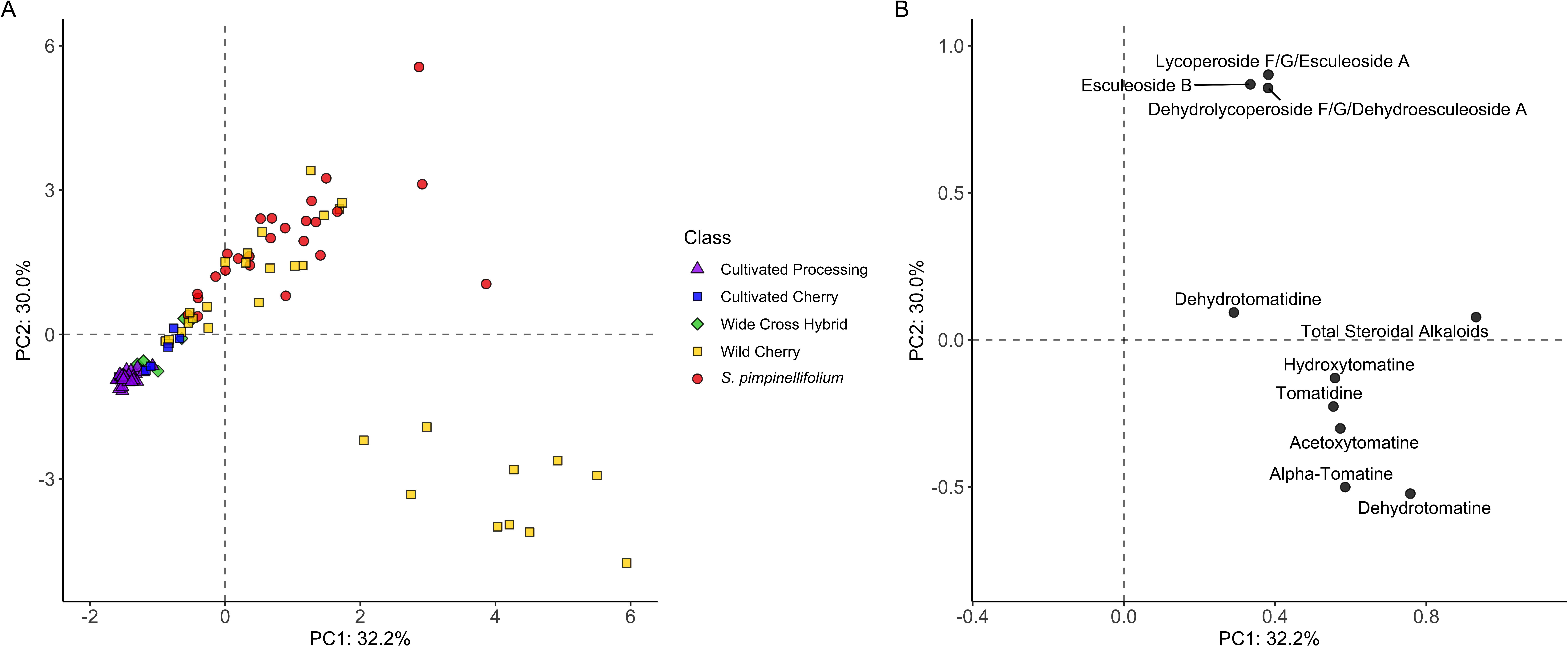
Principal components analysis scores plot (A) and corresponding loading plots (B) for 107 genotypes represented in the diversity panel. Loadings represent vectors of steroidal alkaloids phenotyped in the population and their direction/magnitude indicate their influence on a given principal component. Wild accessions (i.e., wild cherry and *S. pimpinellifolium*) exhibit greater diversity in steroidal alkaloids relative to cultivated accessions. Two distinct subgroups of wild cherry tomatoes appeared to separate based on location in the tomato steroidal alkaloid pathway.

Correlations between metabolites support tradeoffs between of early and late pathway intermediates (Fig. **3**). All steroidal alkaloids associated strongly with adjacent pathway metabolites (e.g., concentrations of lycoperoside F/G/esculeoside A are highly correlated with esculeoside B) and all analytes were positively correlated with total steroidal alkaloids. Steroidal alkaloids with the same mass were also highly, positively correlated with one another when each isomer was considered separately (Fig. **S3**). Negative, but statistically non-significant correlations were observed among some early and late pathway steroidal alkaloids including dehydrotomatine, alpha-tomatine, and acetoxytomatine with dehydrolycoperoside F/G/dehydroesculeoside A, lycoperoside F/G/esculeoside A, and esculeoside B, when considering the full population. To understand how alkaloid correlations differ among tomato type, correlation matrices were generated for each of the five tomato classes. Wild cherry/land-race accessions (Fig. **3c**) exhibited the strongest significant negative correlations among early and late steroidal alkaloid pathway intermediates; consistent with multiple subgroups observable by eye in boxplots (Fig. **1**), PCA (Fig. **2**), and confirmed heuristically using hierarchical clustering (Fig. **S2**).

**Fig. 3.**
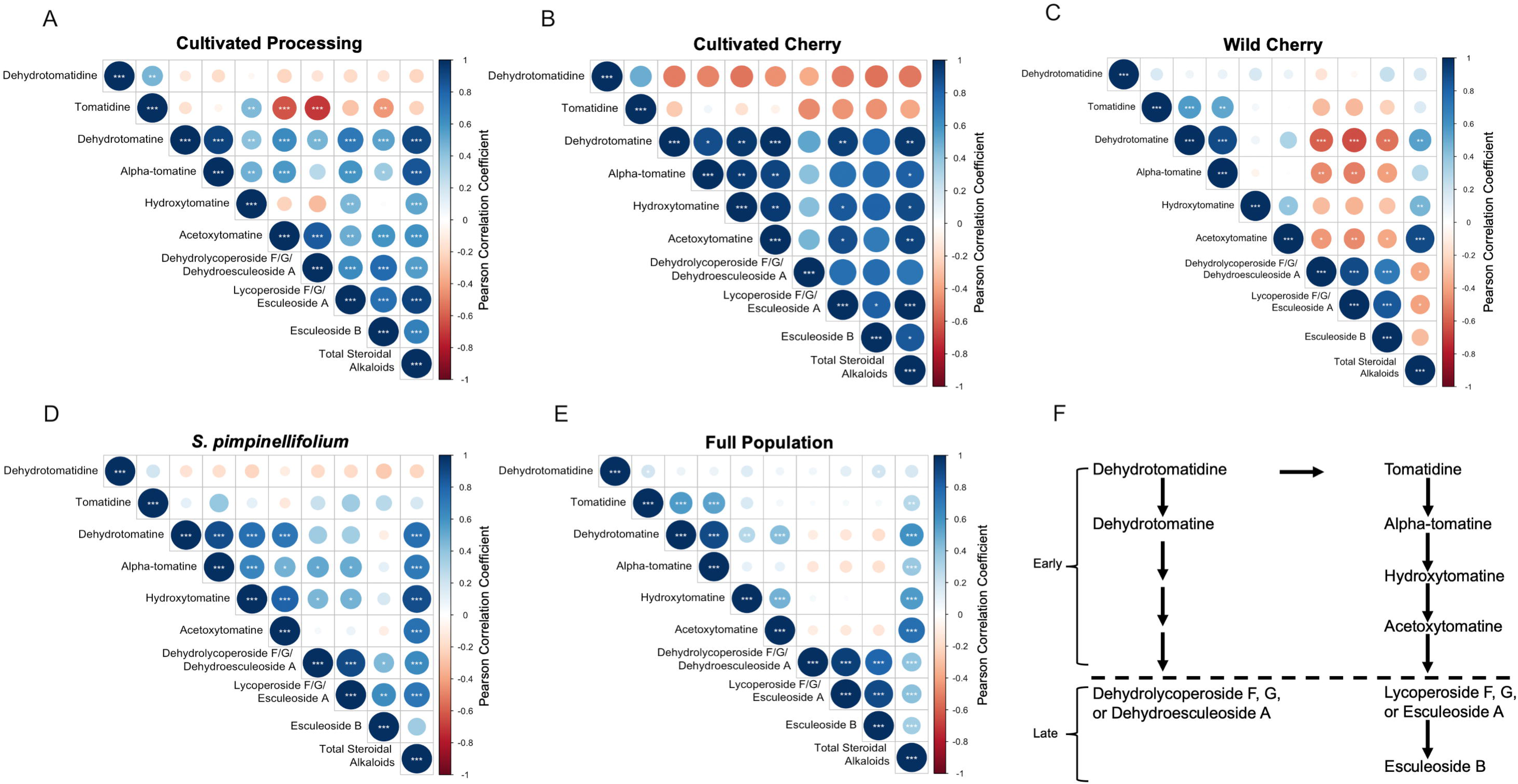
Correlation matrix of all tomato steroidal alkaloids quantified in diversity panel and sub-populations including a) wild cherry, b) cultivated processing, c) cultivated cherry, d) *Solanum pimpinellifolium*, and e) the full diversity panel. Size and darkness of circle indicate intensity of correlation coefficient (see legend on right) and *, **, and *** indicate statistical significance at P<0.05, 0.01, and 0.001, respectively. Cells with no significance indicator were found to be P>0.05. Pathway intermediates tended to correlate strongly with neighboring metabolites in the proposed biosynthetic pathway. All analytes correlated with “total” tomato steroidal alkaloids to varying degrees.

### 1.4.3 Diversity in tomato steroidal alkaloids is under strong genetic control

Multiple growing environments allowed us to partition effects due to genetics, environment, and their interaction on tomato steroidal alkaloid concentrations. Variance partitioning demonstrated that genetic sources of variation were the major contributor to concentrations of most steroidal alkaloids (Table **1**). For alpha-tomatine, hydroxytomatine, acetoxytomatine, and lycoperoside F/G/esculeoside A, broad sense heritability ranged from 0.53 to 0.96 and reliability ranged from 0.46 to 0.68 (Table **1**). Dehydrotomatidine and tomatidine, which represent on average 0.01 and 0.05% of total steroidal alkaloids, respectively, exhibited relatively low proportions of variance explained by “genotype”. The strongest effects were attributed to variation within a single field location. As such, estimates of broad sense heritability and reliability decreased accordingly (Table **1**).

**Table 1.**
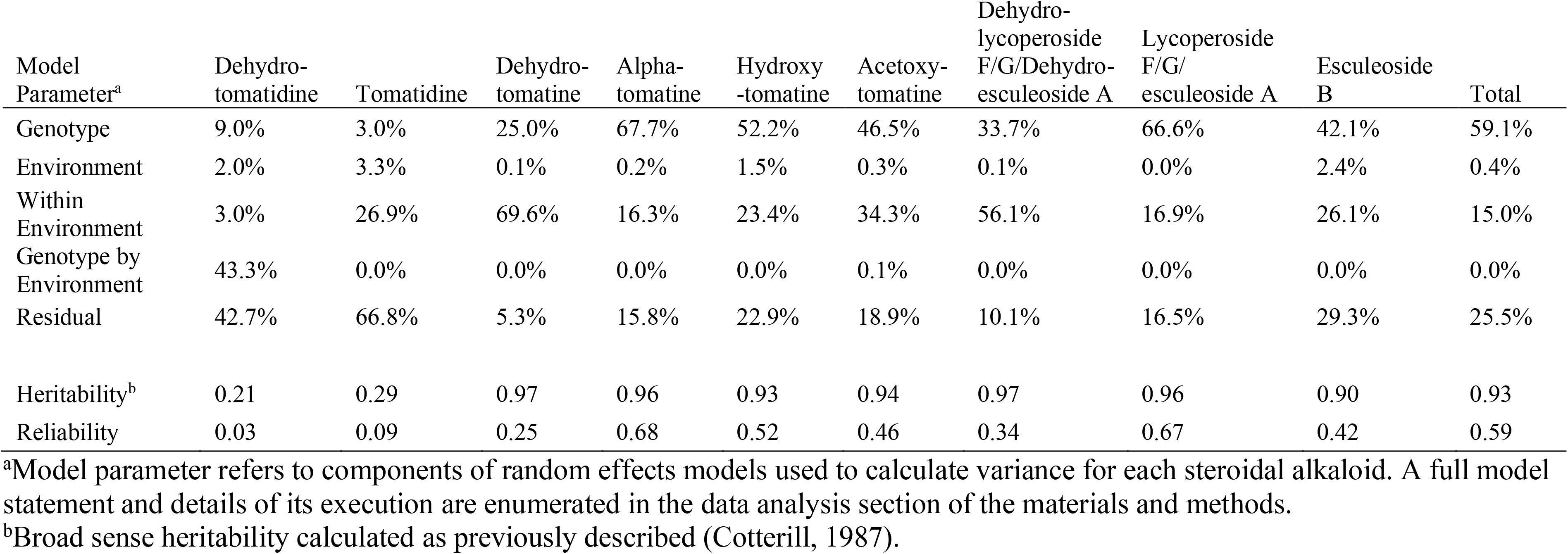
Percentage of total variance due to the contribution of genetics and environment for steroidal alkaloid content.

| Model<br>Parameter<sup>a</sup> | Dehydro-<br>tomatidine | Tomatidine | Dehydro-<br>tomatine | Alpha-tomatine | Hydroxy-<br>tomatine | Acetoxy-<br>tomatine | Dehydro-<br>lycoperoside<br>F/G/Dehydro-<br>esculeoside A | Lycoperoside<br>F/G/<br>esculeoside A | Esculeoside<br>B | Total |
| --- | --- | --- | --- | --- | --- | --- | --- | --- | --- | --- |
| Genotype | 9.0% | 3.0% | 25.0% | 67.7% | 52.2% | 46.5% | 33.7% | 66.6% | 42.1% | 59.1% |
| Environment | 2.0% | 3.3% | 0.1% | 0.2% | 1.5% | 0.3% | 0.1% | 0.0% | 2.4% | 0.4% |
| Within<br>Environment | 3.0% | 26.9% | 69.6% | 16.3% | 23.4% | 34.3% | 56.1% | 16.9% | 26.1% | 15.0% |
| Genotype by<br>Environment | 43.3% | 0.0% | 0.0% | 0.0% | 0.0% | 0.1% | 0.0% | 0.0% | 0.0% | 0.0% |
| Residual | 42.7% | 66.8% | 5.3% | 15.8% | 22.9% | 18.9% | 10.1% | 16.5% | 29.3% | 25.5% |
| Heritability<sup>b</sup> | 0.21 | 0.29 | 0.97 | 0.96 | 0.93 | 0.94 | 0.97 | 0.96 | 0.90 | 0.93 |
| Reliability | 0.03 | 0.09 | 0.25 | 0.68 | 0.52 | 0.46 | 0.34 | 0.67 | 0.42 | 0.59 |

<sup>a</sup>Model parameter refers to components of random effects models used to calculate variance for each steroidal alkaloid. A full model statement and details of its execution are enumerated in the data analysis section of the materials and methods.
<sup>b</sup>Broad sense heritability calculated as previously described (Cotterill, 1987).

To determine the gene action underlying high alpha-tomatine concentration in ripe fruits, LA2213, LA2256, and LA2262 were each crossed to OH8243 (PI 601423; a processing tomato cultivar) and Tainan (PI 647556; a fresh market grape-cherry tomato). F_1_ individuals were not statistically different from the cultivated parent in terms of alpha-tomatine content (Table **S4**). While LA2213, LA2256, and LA2262 were selected for crosses because of their high alpha-tomatine content (Rick *et al*., 1994), other steroidal alkaloids were also profiled in parental and F_1_ individuals. Patterns of inheritance for other steroidal alkaloids varied for each analyte. For example, hydroxytomatine concentrations among cultivated parents and F_1_ individuals were statistically similar for all crosses (Table **S4**). OH8243 and Tainan were numerically higher in the late pathway intermediates dehydrolycoperoside F/G/dehydroesculeoside A and lycoperoside F/G/esculeoside A, though these differences were not statistically significant.

Genome wide association analysis using the QK model, adjusted for structure with different numbers of PC for Q and a kinship matrix (K) were explored. Including a single PC produced the best fit for all models based on BIC. GWAS with a single PC identified 301 putative associations between the ten steroidal alkaloid traits (each individual alkaloid, as well as total steroidal alkaloids) at an FDR-corrected P < 0.05. Phenotypic data, SNP calls, outputs, and summarized results of the GWAS are available in Table **S5-7**. Significant associations were based on 234 unique SNPs spread across all 12 chromosomes. For 278 of the 301 FDR-corrected P<0.05 marker-trait associations, the allele most frequently identified in wild germplasm was associated with an increase in concentration of steroidal alkaloids.

Fig. **4** displays Manhattan plots of steroidal alkaloids, highlighting different genetic associations among early and late pathway intermediates and condensed the large number of single-marker trait associations into a more defined set of loci based on linkage. Thirty-six associations for alpha-tomatine reduce to 10 loci on 9 chromosomes. Associations for early pathway intermediates tended to be stronger and more frequent than those of late pathway intermediates. For example, loci were detected by multiple linked markers for alpha-tomatine on 8 of 12 chromosomes while only four loci were identified for dehydrolycoperoside F/G/dehydroesculeoside A on chromosome 1, 9 and 12. Markers associated with multiple steroidal alkaloids tended to be associated with neighboring metabolites (e.g., the association of solcap_snp_sl_100848 with hydroxytomatine and acetoxytomatine).

**Fig. 4.**
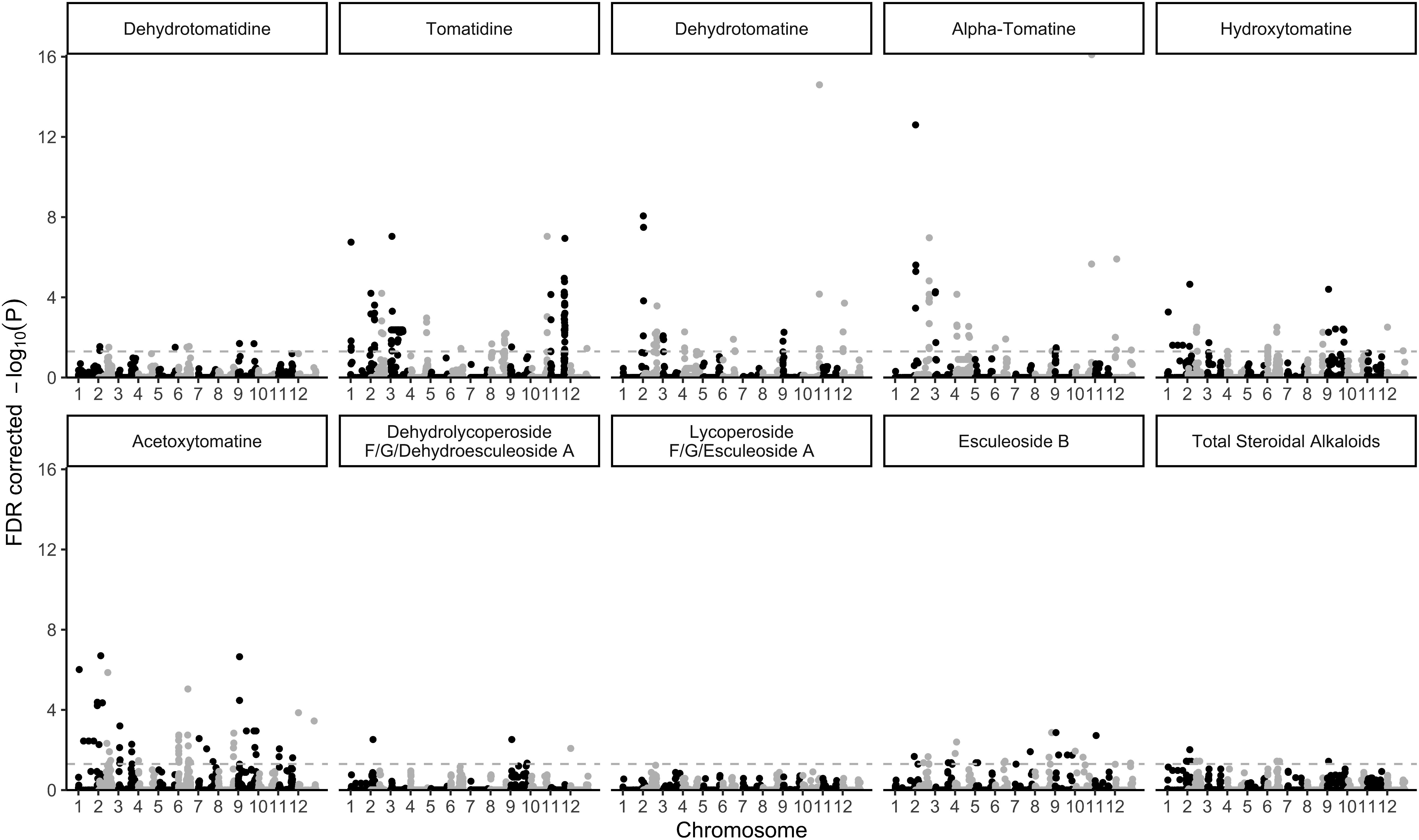
Manhattan plots of steroidal alkaloids used in GWAS. The dashed line indicates SNPs above a significance threshold of −log_10_ Benjamini-Hochberg corrected P-values of 1.3 (P<0.05).

### 1.4.4 Association determined by GWAS are validated in a biparental population

The BC_1_S_1_ material generated from crossing OH8243 (cultivated processing; low alpha-tomatine in ripe fruit) with LA2213 (land-race cherry; high alpha-tomatine in ripe fruit) provided a separate, independent population to confirm metabolic QTL (mQTL) found in the GWAS. Phenotypic data, SNP calls, and summarized results are available in Table **S8,9**. Analysis of backcross progeny confirmed a role for a locus on chromosome 3 in controlling multiple early pathway steroidal alkaloids as well as total concentration in tomato (Table **2**). These associations are also visualized with Manhattan plots seen in Fig. **5**. SNPs on chromosome 3 (solcap_snp_sl_7942, solcap_snp_sl_7939, solcap_snp_sl_7919, solcap_snp_sl_5761, and solcap_snp_sl_5656) were strongly associated with tomato steroidal alkaloids from the first half of the pathway (dehydrotomatine, alpha-tomatine, hydroxytomatine, and acetoxytomatine), as well as total tomato steroidal alkaloids. Later pathway metabolites (dehydrolycoperoside F/G/dehydroesculeoside A, lycoperoside F/G/esculeoside A, and esculeoside B) did not exhibit this association with loci on chromosome 3. Markers on chromosomes 10 (solcap_snp_sl_13202 and solcap_snp_sl_46386) and 11 (SL10890_654) were associated with the late pathway steroidal alkaloid lycoperoside F/G/esculeoside A.

**Table 2.**
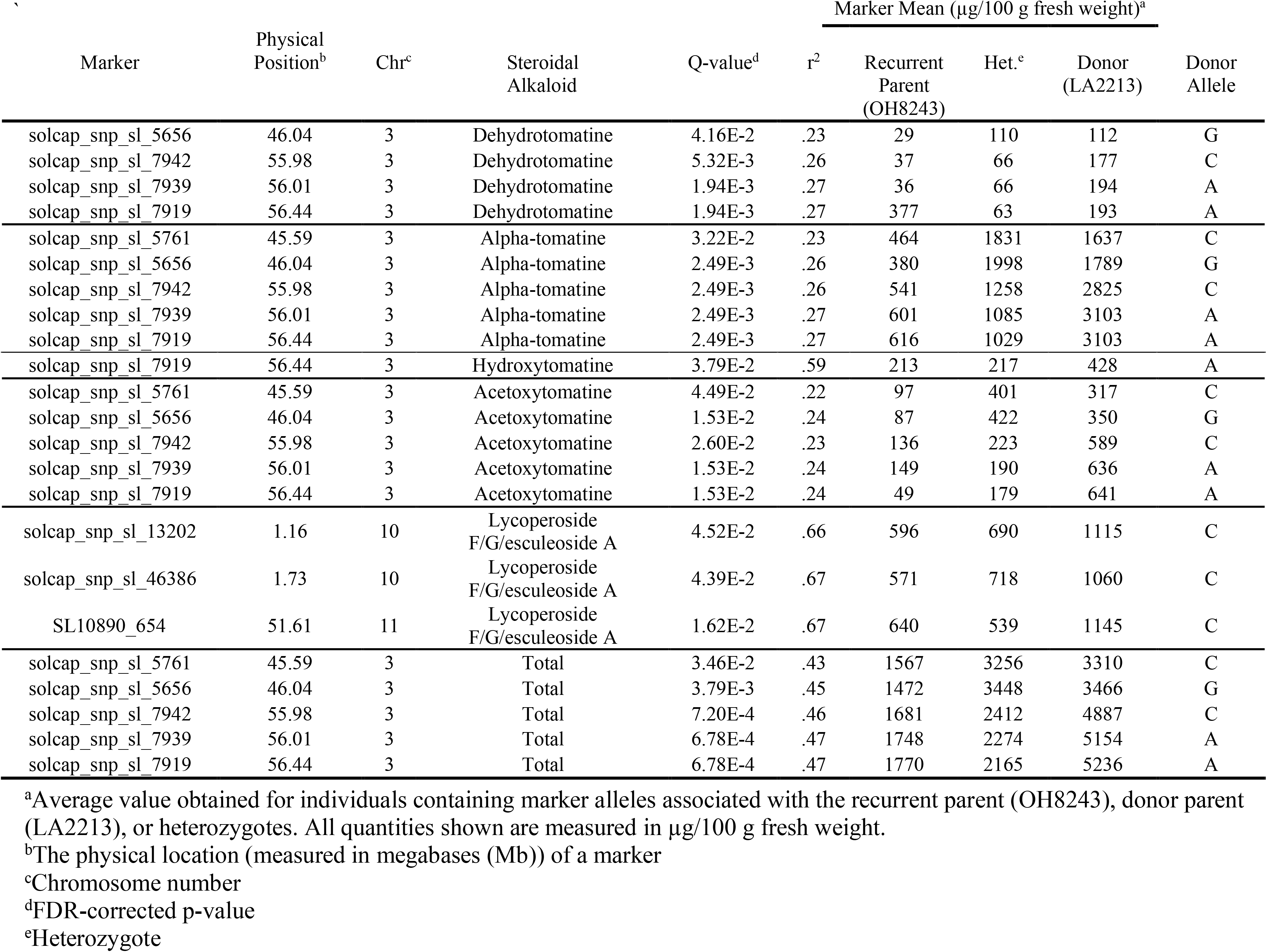
Markers associated (FDR-corrected P <0.05) with tomato steroidal alkaloids in the BC_1_S_1_ validation population. Marker means reported in μg/100 g fresh weight.

**Fig. 5.**
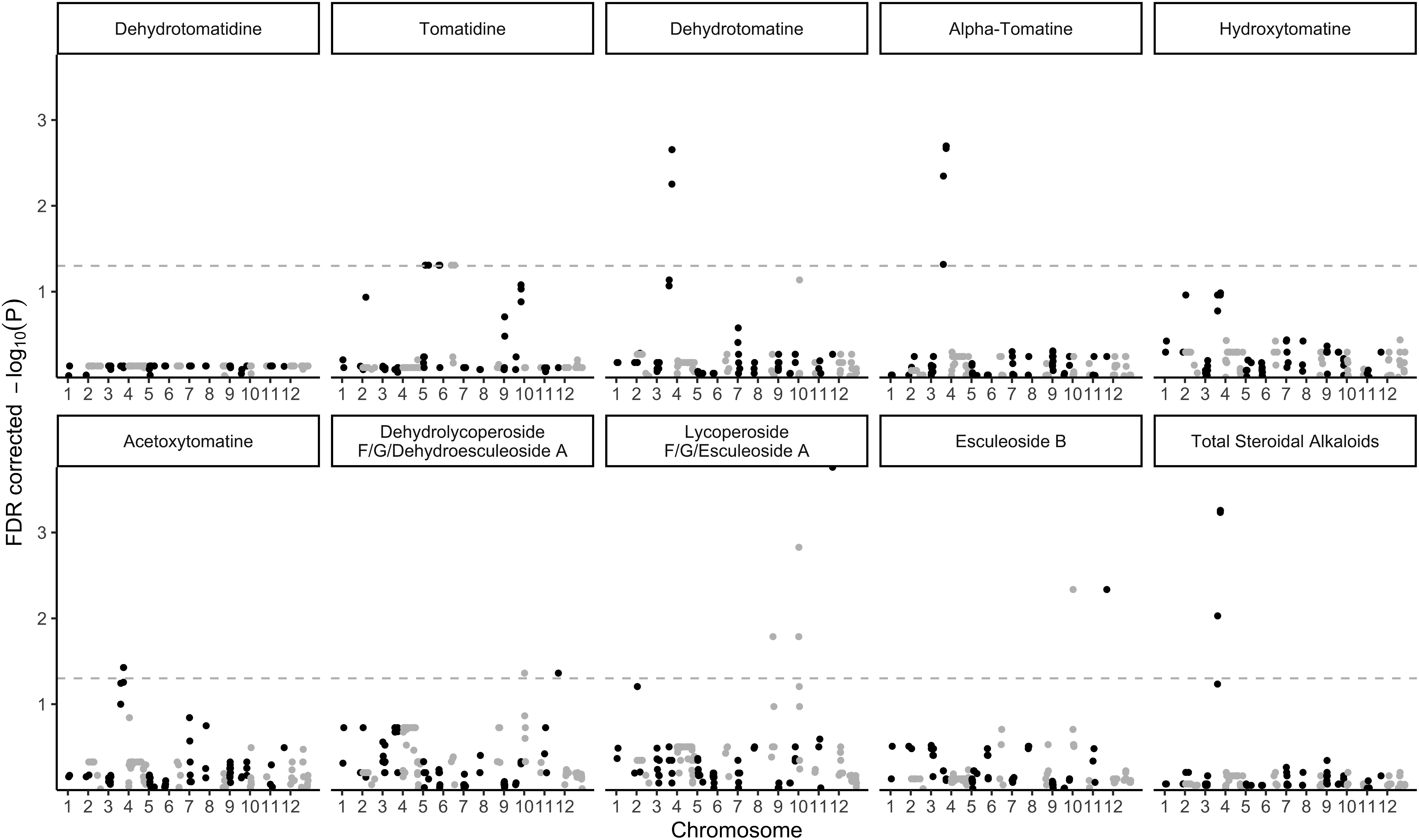
Manhattan plots of steroidal alkaloids used in QTL analysis on the BC_1_S_1_ validation population. The dashed line indicates SNPs above a significance threshold of −log_10_ Benjamini-Hochberg corrected P-values of 1.3 (P<0.05).

## 1.5 Discussion

### 1.5.1 Diversity for steroidal alkaloids is greatest in land-race and wild species

Steroidal alkaloids were found in higher concentrations in land-race and wild accessions compared to cultivated cherry and processing classes (Fig. **1**). Alpha-tomatine was a minor component of steroidal alkaloid profiles except for cultivated processing and wild cherry material; the latter being inflated due to five high alpha-tomatine land-races (LA1654, LA2213, LA2256, LA2262, and LA2308; Table **S1,2**). This finding agrees with the current model that early pathway intermediates are converted to late pathway during ripening (Yamanaka *et al*., 2009; Itkin *et al*., 2011). Average values for individual alkaloids ranged between 0.1 and 7.6 mg/100 g fresh weight (Table **S1,2**), demonstrating tomato steroidal alkaloids exist in concentrations higher than previously thought (Friedman, 2002; Caprioli *et al*., 2014; Baldina *et al*., 2016), and consistent with more recent reports (Steinert *et al*., 2020). Prominent tomato phytochemical classes like carotenoids are often found between 0.09 to 9.93 mg/100 g fresh weight (Dzakovich *et al*., 2019), suggesting steroidal alkaloids can be found in comparable concentrations. Exceptional levels were observed. For example, lycoperoside F/G/esculeoside A were quantified in concentrations of up to 60 mg/100g fresh weight in wild accessions (PI-155372; Table **S1, 2**), suggesting substantial diversity for concentrations of steroidal alkaloids in wild tomato. Lower average concentrations and diversity of steroidal alkaloids in cultivated processing material fit with the hypothesis that steroidal alkaloids were under strong negative selection during domestication (Zhu *et al*., 2018).

### 1.5.2 Land-race and wild cherry accessions can be differentiated by early or late pathway steroidal alkaloid concentrations

A noticeable bifurcation in PCA scores plot of the diversity panel alkaloid concentrations can be seen in Fig. **2a**. Separation on PC1 is driven by total steroidal alkaloid concentrations, thus differentiating between cultivated and land-race and wild accessions. Individuals separated on PC2 appeared to be distinguished based on the abundance of specific steroidal alkaloids. The loadings plot in Fig. **2b** visualizes the separation of early and late steroidal alkaloid biosynthetic pathway intermediates and confirms that these profiles are driving the separation along PC2 (Fig. **2a**). Ten S. *lycopersicum* var. *cerasiforme* individuals appear separated in the PCA scores plot from other *S. lycopersicum* var. *cerasiforme* and *S. pimpinellifolium* accessions, and this relationship was confirmed heuristically by hierarchical clustering analysis (Fig. **S2**). These *S*. *lycopersicum* var. *cerasiforme* individuals include the high alpha tomatine accessions LA2213, LA2256, and LA2262 previously described (Rick *et al*., 1994) as well as several other accessions from the same region of Peru. Correlation analyses among tomato steroidal alkaloids reflect a differential pattern of pathway intermediate accumulation among germplasm classes as well (Fig. **3**). Late-pathway tomato steroidal alkaloids tend to be negatively correlated with early-pathway metabolites in the land-race and wild cherry accessions but not other germplasm groups (Fig. **3c**). This finding suggests differential regulation of early and late parts of the pathway in some land-race and wild cherry selections. Exploring tomato germplasm within the red-fruited clade may therefore offer insight into the genetic regulation of steroidal alkaloids.

### 1.5.3 Tomato steroidal alkaloids are heritable

Heritability and reliability estimates demonstrate that production of tomato steroidal alkaloids such as alpha-tomatine, hydroxytomatine, acetoxytomatine, and lycoperoside F/G/esculeoside A are under strong genetic control (Table **1**). Two previous reports provide estimates of broad sense heritability between 0.35 and 0.95, with most being above 0.50 (Alseekh *et al*., 2015; Zhu *et al*., 2018). These data are consistent with our findings. All crosses between high alpha-tomatine land-race tomatoes (LA2213, LA2256, and LA2262) with low alpha-tomatine contemporary cultivated tomatoes show that low alpha-tomatine in ripe fruits is a dominant trait (Table **S4**). The typical pattern in contemporary cultivated tomato is for the early pathway metabolites such as alpha-tomatine to be converted to late-pathway intermediates such as lycoperoside F/G/esculeoside A during ripening (Iijima *et al*., 2009; Cárdenas *et al*., 2019; Szymański *et al*., 2020). The recessive nature of high alpha-tomatine suggests that elevated levels in ripe fruit occur due to a loss of function and a block in the shift from early to late pathway steroidal alkaloids. These results are consistent with previous findings where LA2213, LA2256, and LA2262 were initially characterized and crossed with low alpha-tomatine varieties (Rick *et al*., 1994). The trade-off between late and early pathway metabolites and strong heritability are consistent with changes in pathway regulation.

### 1.5.4 A QTL on chromosome 3 controls multiple early pathway steroidal alkaloids

GWAS illuminated hundreds of putative associations between molecular markers and steroidal alkaloids with 85% of these associations occurring with tomatidine, dehydrotomatine, alpha-tomatine, hydroxytomatine, and acetoxytomatine. These analytes appear to be driving the separation visualized along PC2 (Fig. **2**). Associations on chromosome 3 for dehydrotomatine, tomatine, hydroxytomatine, acetoxytomatine, and total steroidal alkaloids were validated in a biparental backcross population. Associations for the late pathway intermediate lycoperoside F/G/esculeoside A detected in the bi-parental backcross population on chromosomes 10 and 11 were not observed in the diversity panel GWAS. The identification of QTL associated with the concentration of early metabolites on chromosome 3 (Table 2, Fig. **5**.) is a novel finding and drives the separation of germplasm based on metabolite loading (Fig. **2b**). These data suggest that the coordinate regulation of early and late tomato steroidal alkaloid pathway steps is under distinct genetic control.

Previous studies seeking to map genes involved in the biosynthesis of tomato steroidal alkaloids have used wild tomato species with an emphasis on *S. pennellii* LA0716. These studies have found QTL on all chromosomes of the tomato genome (Alseekh *et al*., 2015), but the majority of reports highlight chromosomes 2, 3, 7, 10, and 12 (Itkin *et al*., 2013; Baldina *et al*., 2016; Ballester *et al*., 2016; Zhu *et al*., 2018; Szymański *et al*., 2020). The GAME genes have been found on chromosomes 1, 2, 7, and 12 (Itkin *et al*., 2013). A previous mQTL study identified multiple candidate genes in a sweep region at the end of chromosome 3 (Zhu *et al*., 2018). Among these candidates were two ethylene-responsive transcription factors. Many of the steps in the steroidal alkaloid biosynthetic pathway that have been elucidated are modulated by hormones, such as ethylene, during ripening (Iijima *et al*., 2008, 2009; Itkin *et al*., 2011). A transcription factor that can affect multiple steps in the biosynthetic pathway may be responsible given that multiple steroidal alkaloids in our population appear to be affected by the same QTL on chromosome 3. This QTL has not been reported before and may be unique to the germplasm we selected.

Quantitative profiling of red-fruited species demonstrates that tomato steroidal alkaloids are an abundant and diverse class of secondary metabolites in ripe fruit. Much of the focus on elucidating the genetic architecture of these compounds has been within the context of their potential bitterness or perceived negative effects on human health (Rick *et al*., 1994; Itkin *et al*., 2013; Cárdenas *et al*., 2016, 2019; Ballester *et al*., 2016), despite numerous reports indicating that steroidal alkaloids may be associated with positive health outcomes (Cayen, 1971; Lee *et al*., 2004; Choi *et al*., 2012b; Dyle *et al*., 2014), and accumulate in the body after consumption (Cichon *et al*., 2017; Cooperstone *et al*., 2017; Hövelmann *et al*., 2019, 2020). By purposefully creating germplasm to study the control of steroidal alkaloid biosynthesis in the red-fruited clade, we generated quantitative evidence of a metabolic tradeoff between early and late pathway intermediates. This pattern was observable through phenotypic data, unsupervised learning, correlation among pathway intermediates, and genetic analysis. Our results confirm that the previous model hypothesizing a degradation of tomato steroidal alkaloids during ripening (Rick *et al*., 1994), does not accurately reflect the shift in metabolites that occurs in all of the red-fruited clade. We identified a previously unreported QTL on chromosome 3 associated with the accumulation of dehydrotomatine, alpha-tomatine, hydroxytomatine, acetoxytomatine, and total steroidal alkaloids in ripe tomato fruits. This association was derived from wild cherry (including land-race) accessions from San Martin and La Libertad regions of Peru and strongly supported in the biparental population derived from LA2213. Fine mapping experiments to identify gene(s) underlying this QTL will allow for the exploitation of steroidal alkaloids to better understand their role in defense, flavor, and human health.

## Acknowledgments

We thank the crew at the North Central Agriculture Research Station of OSU, particular Matt Hoffelich and Frank Thayer for excellent tomato care. We would also like to thank Troy Aldrich and Jiheun Cho for helping coordinate planting and seed saving.

Financial support for this work was provided by an Ohio Agricultural Research and Development Center Early Career Investigator Award, USDA National Needs Fellowship (2014-38420-21844) and Hatch funds (OHO01470, OHO01405), Foundation for Food and Agricultural Research New Innovator Award, and Foods for Health, a focus area of the Discovery Themes at OSU.

The authors have no conflicts of interest to disclose.

## Supporting Information Figure and Table Legends

Fig. **S1**. Map of Central and South America indicating collection location of wild germplasm.

Fig. **S2**. Principal components analysis scores plot of wild cherry tomato accessions overlaid with hierarchical clustering assignments.

Fig. **S3**. Correlation plot of all steroidal alkaloids and isomers measured in this study.

Table **S1**. Raw data and metadata of the diversity germplasm. Data are presented in μg/100g FW

Table **S2**. Genotypic means and standard deviations of each steroidal alkaloid quantified.

Table **S3**. Genotypic least squares means of each steroidal alkaloid quantified.

Table **S4**. Means and standard deviations of each steroidal alkaloid of parental and F_1_ material. Letters represent significance groups as determined by a Tukey-Kramer post-hoc test.

Table **S5**. Genotypic means of each steroidal alkaloid and SNP calls for diversity panel germplasm members that were previously genotyped using the SolCAP Illumina Infinium Array (7,720 SNPs).

Table **S6**. Marker-trait associations generated by GWAS of the diversity panel with a minor allele frequency cutoff of 10%

Table **S7**. Summary of significant marker trait associations generated by GWAS of the tomato diversity panel.

Table **S8**. Genotypic means of each steroidal alkaloid and SNP calls for the BC_1_S_1_ biparental mapping population using the 384 marker SolCAP array.

Table **S9**. Marker-trait associations generated by QTL analysis of the BC1S1 validation population with a minor allele frequency cutoff of 10%

