## Supplementary figures and images for "Biosynthesis of Steroidal Alkaloids Are Coordinately Regulated and Differ Among Tomatoes in the Red-Fruited Clade"

### Supplemental Figure 2

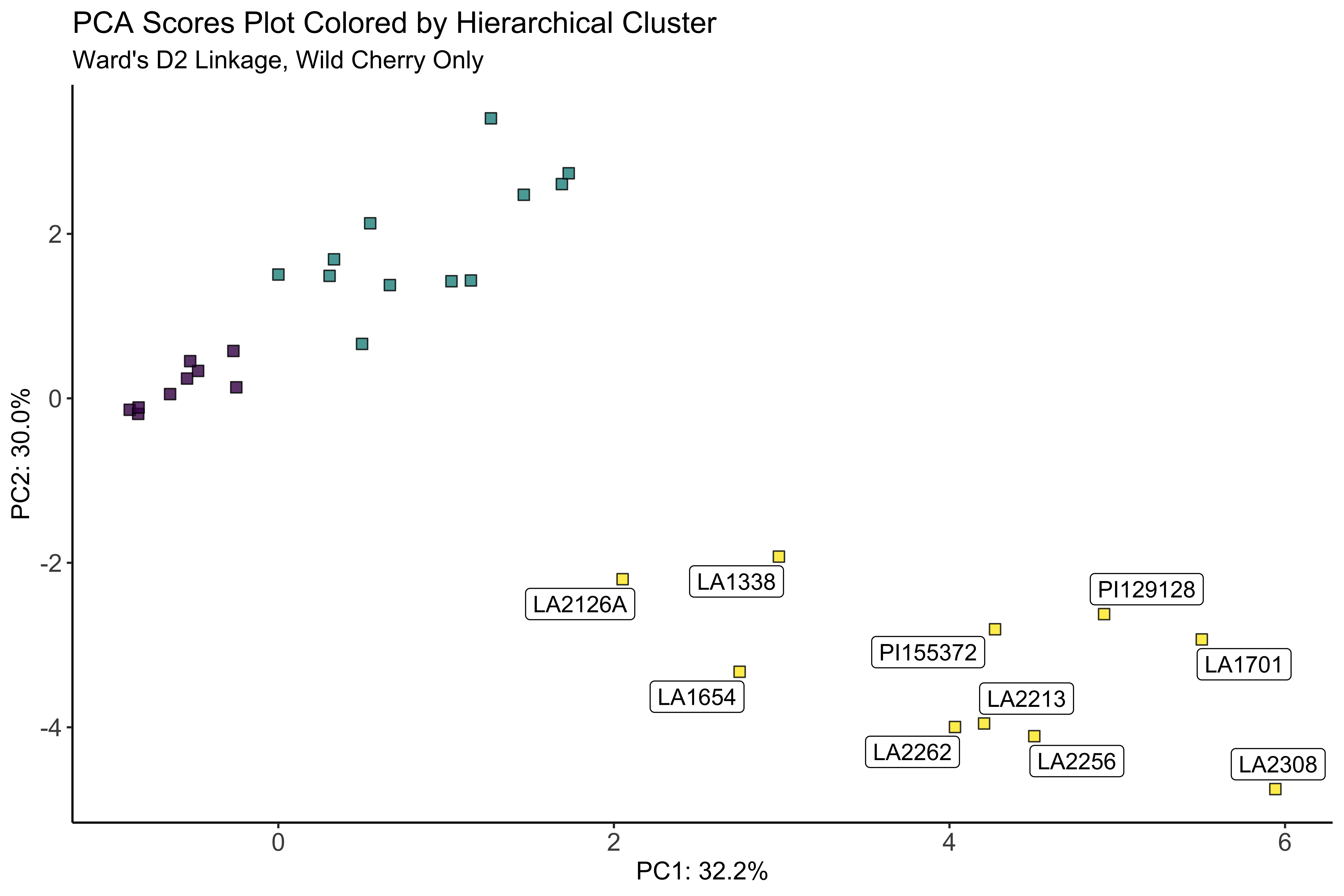

### Supplemental Figure 3

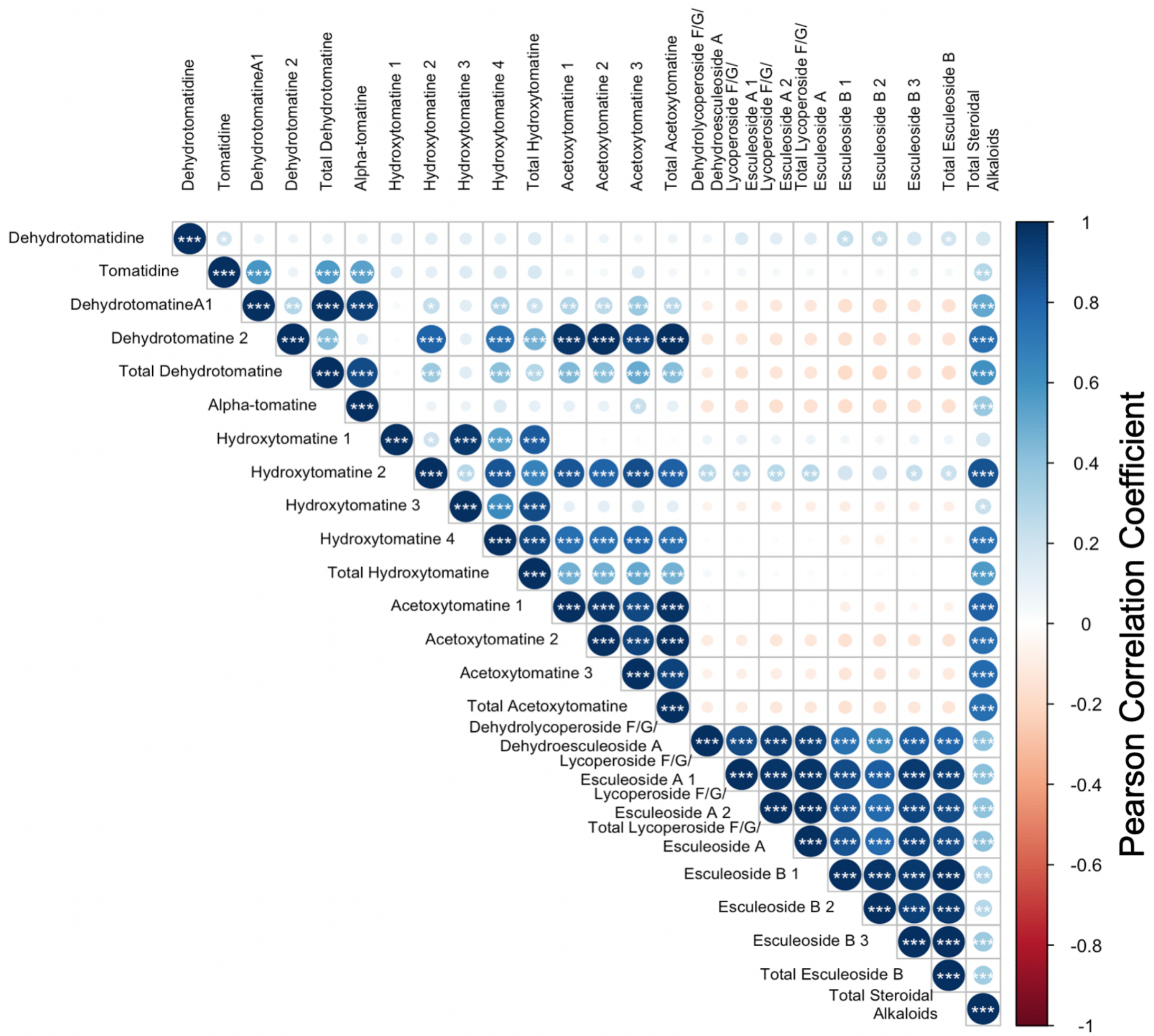

### Supplementary Figure 1

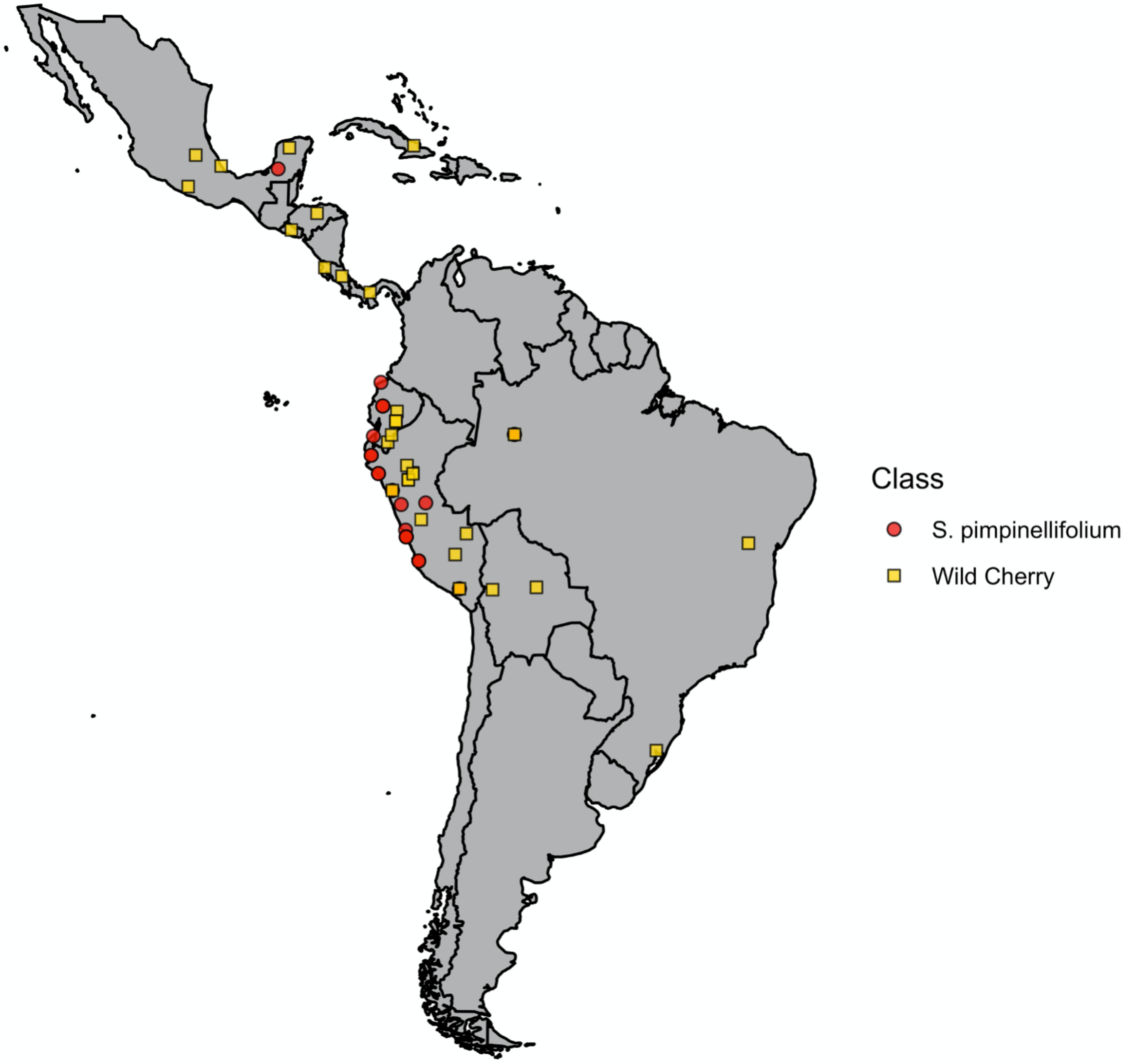
